# Structural and functional elucidation of NF-κB signaling by the p75 neurotrophin receptor through recruitment of TRADD

**DOI:** 10.1101/2021.02.07.430167

**Authors:** Ning Zhang, Lilian Kisiswa, Ajeena Ramanujan, Zhen Li, Eunice Weiling Sim, Wensu Yuan, Carlos F. Ibáñez, Zhi Lin

## Abstract

p75 neurotrophin receptor (p75^NTR^) is a critical mediator of neuronal death and tissue remodeling and has been implicated in various neurodegenerative diseases. The death domain (DD) of p75^NTR^ is an intracellular signaling hub and has been shown to interact with diverse adaptor proteins. However, the structural mechanism and physiological relevance of the adaptor protein TRADD in neuronal p75^NTR^ signaling remain poorly understood. Here we report an NMR structure of the complex between p75^NTR^-DD and TRADD-DD and elucidate the structural basis of specific DD recognition in the p75^NTR^/TRADD signaling pathway. Furthermore, we identify spatiotemporal overlap of p75^NTR^ and TRADD expression in developing cerebellar granule neurons (CGNs) at early postnatal stages and reveal the functional role of TRADD recruitment to p75^NTR^ in the regulation of canonical NF-κB signaling and cell survival in CGNs. Our results provide a new structural framework for understanding how the recruitment of TRADD to p75^NTR^ through DD interactions creates a membrane-proximal platform to propagate downstream signaling in developing neurons.

## 1. Introduction

p75 neurotrophin receptor (p75^NTR^), also known as tumor necrosis factor receptor superfamily (TNFRSF) 16 and nerve growth factor receptor (NGFR), is the first receptor identified for a family of neurotrophic growth factors known as the neurotrophins (NTs).^[1]^ p75^NTR^ is a critical mediator of neuronal death and tissue remodeling and has been implicated in a variety of neurodegenerative diseases.^[2]^ In the early development of mammals, p75^NTR^ is highly expressed in the nervous system and plays an essential role in promoting nerve growth. Its expression is significantly downregulated in the adult and only observed in a small number of neuronal subpopulations; however, lesions to the nervous system strongly stimulate p75^NTR^ re-expression.^[2]–[3]^ In many cancers, such as melanoma and breast cancer, p75^NTR^ is also expressed as a tumor promoter or suppressor, depending on cancer types.^[4]^ Similar to other TNFRSF members, p75^NTR^ does not have intrinsic catalytic activity, and its signaling depends on the recruitment of intracellular interactors.^[5]^

p75^NTR^ consists of an extracellular cysteine-rich domain (ECD), a single-pass transmembrane domain (TMD) involved in the formation of an intermolecular disulfide bridge, and an intracellular domain (ICD), containing an unstructured juxtamembrane domain (JMD) and a helical death domain (DD).^[5]–[6]^ Upon binding extracellular signals, such as the neurotrophin nerve growth factor (NGF), the p75^NTR^-ECD undergoes significant conformational changes from an open to a closed state.^[7]^ This movement further propagates to the p75^NTR^-DD through the disulfide-bonded p75^NTR^-TMD, leading to separation of the p75^NTR^-DD homodimer and exposure of active sites on the p75^NTR^-DD surface for recruitment of DD interactors.^[8]^ As a signaling hub, the p75^NTR^-DD has been shown to interact with various intracellular molecules to regulate different signaling cascades, including NF-κB, RhoA, and JNK/caspase pathways in diverse cellular contexts.^[5, 9]^ Among the many proteins that interact with the p75^NTR^-ICD, TRADD (TNF receptor associated death domain) is rather unique due to its ability to create a platform on the membrane for recruitment of additional proteins for downstream signaling.^[10]^ TRADD is a 34 kDa protein with two functional domains (N- and C-terminal domains) connected by an unstructured peptide of ~38 amino acids.^[11]^ The C-terminal domain of TRADD contains a novel DD. TRADD was initially identified as an adaptor protein for tumor necrosis factor receptor 1 (TNFR1) signaling since its DD can interact with the TNFR1-DD in response to TNF activation.^[10], [12]^ It has also been reported that TRADD interacts with p75^NTR^ and promote NGF-dependent signaling that opposes cell death in breast cancer cells.^[13]^ Aside from that first report, there are no other studies on the role and physiological relevance of TRADD in p75^NTR^ signaling. In particular, it remains unknown whether and, if so, how TRADD participates in p75^NTR^ signaling in neurons. Here, we report the NMR complex structures of p75^NTR^-DD:TRADD-DD and reveal the specific mechanism of TRADD-DD recognition by the p75^NTR^-DD. We also identify the functional role of TRADD recruitment to p75^NTR^ through DD interaction in the regulation of NF-κB signaling and cell survival in developing cerebellar granule neurons (CGNs), a physiologically relevant neuronal subpopulation which development is strongly regulated by p75^NTR^ signaling.^[9b, 14]^

## 2. Results and Discussion

### 2.1 Solution structure of the complex between the p75NTR-DD and the TRADD-DD

Our recent NMR structural studies have identified a direct binding of the TRADD-DD to the p75^NTR^-DD *in vitro* and characterized a novel structural fold of the TRADD-DD in the DD superfamily.^[11b]^ The TRADD-DD contains a unique β-hairpin motif, which is not found in any other known DDs. However, it was unclear if this unusual structural element has a specific role in DD-DD interactions. To better understand the structural mechanisms of DD recognition in the p75^NTR^/TRADD interaction, we first determined the solution structure of the complex between the p75^NTR^-DD and the TRADD-DD by multidimensional nuclear magnetic resonance (NMR) spectroscopy (Table 1). Due to the fully and partially exposed hydrophobic residues on the surface, the TRADD-DD exhibits aggregation-prone property in solution. Thus, the protein complex of the p75^NTR^-DD and TRADD-DD for NMR structural studies was prepared in salt-free water, where TRADD-DD aggregation can be minimized and the protein still retains its function for binding to the p75^NTR^-DD (Figure S1, Supporting Information).^[11b, 15]^ The ensemble of ten lowest-energy structures of the p75^NTR^-DD:TRADD-DD complex and a representative cartoon structure are depicted in Figure 1A and B, respectively. Multiple refinements converged to a mean backbone root mean square deviation (RMSD) of 0.32 ± 0.09 Å in the structural region of the complex (G334-E420 of the p75^NTR^-DD and T201-L302 of the TRADD-DD). The representative slice of the detected intermolecular nuclear Overhauser effects (NOEs) shows that the residue K349 from the p75^NTR^-DD is critical for binding the TRADD-DD (Figure 1C). The structure of the p75^NTR^-DD in the complex was close to that in the p75^NTR^-DD homodimer with a pairwise RMSD of ~1.1 Å. A small observable movement of helix orientation occurs in helixes α1 and α4 (Figure S2, Supporting Information). In comparison, the TRADD-DD in the complex is nearly the same as TRADD-DD monomer with a pairwise RMSD of ~0.5 Å. Therefore, the recruitment of TRADD to p75^NTR^ does not impose noticeable conformational changes on either of these death domains.

**Table 1.** NMR and refinement statistics for the complex between the p75^NTR^-DD and the TRADD-DD.

| Parameters |  |
| --- | --- |
| NMR distance & dihedral constraints |  |
| Distance constraints |  |
| Total NOE | 4651 |
| Intra-residue | 1535 |
| Inter-residue |  |
| Sequential ($\vert i-j\vert = 1$) | 1157 |
| Medium-range ($\vert i-j\vert \leq 4$) | 1058 |
| Long-range ($\vert i-j\vert \geq 5$) | 901 |
| Intermolecular NOE | 39 |
| Total dihedral angle restraints <sup>a)</sup> | 320 |
| Structure Statistics |  |
| Violations (mean and s.d.) |  |
| Distance constraints (Å) | 0.38±0.00 |
| Dihedral angle constraints (°) | 4.34±0.38 |
| Max. dihedral angle violation (°) | 4.98 |
| Max. distance constraint violation (Å) | 0.39 |
| Ramachandran Plot <sup>b)</sup> |  |
| Most favoured regions | 91.4% |
| Additional allowed regions | 7.9% |
| Generously allowed regions | 0.7% |
| Disallowed regions | 0.0% |
| Average RMSD (Å) <sup>c)</sup> |  |
| Backbone atoms | 0.32±0.09 |
| Heavy atoms | 0.70±0.06 |

<sup>a)</sup> Dihedral angle constraints were generated by TALOS based on $C_\alpha$ , $C_\beta$ , $H_\alpha$ and N chemical shifts. <sup>b)</sup> The selected residues are 334-420 of p75<sup>NTR</sup>-DD and 201-302 of TRADD-DD in the structural region. <sup>c)</sup> Average r.m.s. deviation (RMSD) to the mean structure was calculated among 10 refined structures. Superimposed residues are 334-420 of p75<sup>NTR</sup>-DD and 201-302 of TRADD-DD. The total AMBER energy is -9014±47 kcal/mol.

**Figure 1.**
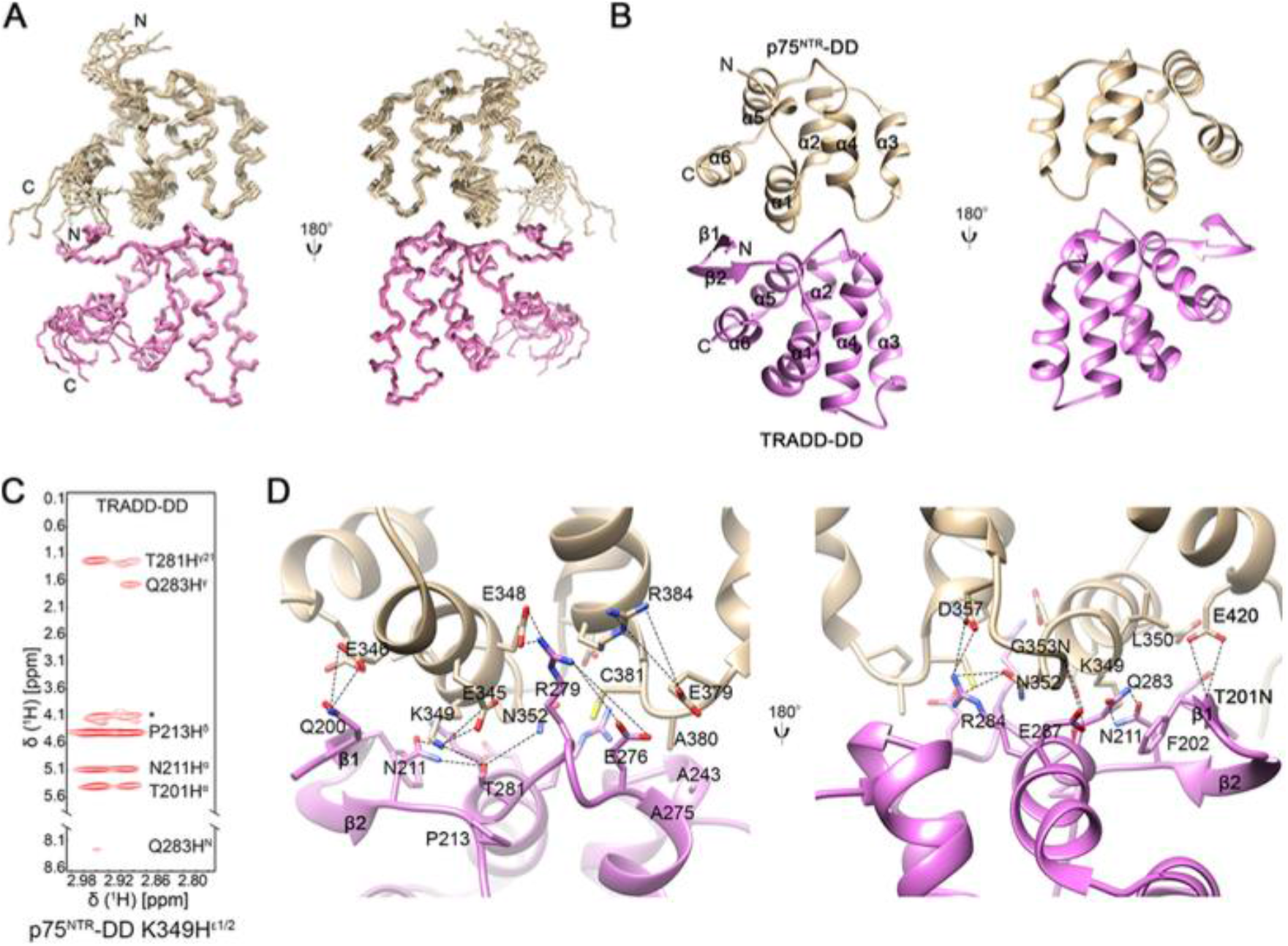
Solution structure of p75^NTR^-DD:TRADD-DD complex. A) Superposition of backbone heavy atoms of the 10 lowest-energy complex structures of the p75NTR-DD:TRADD-DD. N- and C-termini of DDs are indicated. B) Ribbon drawing of p75^NTR^-DD:TRADD-DD. C) Representative slice from the ^13^C,^15^N-filtered 3D NOESY spectrum. *, ambiguous NOE peaks. The p75^NTR^-DD was labeled with ^13^C and ^15^N, and the TRADD-DD was unlabeled. D) Detail of binding interface in the p75^NTR^-DD:TRADD-DD complex. Key residues at the binding interface are labelled and depicted as stick models. Close distances between nitrogen and oxygen atoms (~5Å or less) are showed in dash lines.

#### DD interaction interface

High resolution of the NMR structure allows us to closely inspect the binding interface in the complex. Figure 1D shows that the DD interaction interface is mainly formed by helices α1, α6, and α3-α4 loop of the p75^NTR^-DD and β-hairpin (β1 and β2), helices α5, α2-α3 and α4-α5 loops of the TRADD-DD. Hydrophobic interactions between residues L350 and A380 of the p75^NTR^-DD and F202, A243, and A275 of the TRADD-DD also contribute to the binding of two DDs. We have previously shown that the β-hairpin motif is indispensable for the TRADD-DD to fold correctly in water solution.^[11b]^ The interactions observed in the p75^NTR^-DD:TRADD-DD complex highlight the structural importance of this β-hairpin motif in complex assembly between DDs. Electrostatic interactions were also found to play a crucial role in the binding interface, and an extensive set of charged and polar residues, including Arg, Lys, Glu, Asp, Gln, Asn, were identified (Figure 1D). Hydrogen bonding between E348 in the p75^NTR^-DD and R279 in the TRADD-DD as well as between N352 in the p75^NTR^-DD and R284 in the TRADD-DD was detected. Importantly, isothermal titration calorimetry (ITC) assays as well as co-immunoprecipitation experiments verified the functional relevance of six key interface residues, namely E345, K349, and E379 in the p75^NTR^-DD and E276, R279, and R284 in the TRADD-DD. Single-point mutation of these charged residues to Ala significantly diminished TRADD interaction with p75^NTR^ in ITC assays using purified proteins and in transfected HEK 293T cells (Figure 2, Figure S3 and S4, Supporting Information). One of the most critical residues in the binding interface is K349 from the p75^NTR^-DD, which shows a number of interactions with residues N211, T281, Q283, and P213 from the TRADD-DD. A single K349A mutation can nearly completely disrupt the interaction between the TRADD-DD and the p75^NTR^-DD, leading to undetectable binding affinity measured by ITC assay (Fig 2B). We further confirmed this data in transfected HEK cells where the interaction between WT TRADD-DD and p75^NTR^-DD in the K349A mutation was undetectable (Fig 2C). Taken together, these mutagenesis studies highlight the critical contribution of the charged surface to DD interaction between p75^NTR^ and TRADD.

**Figure 2.**
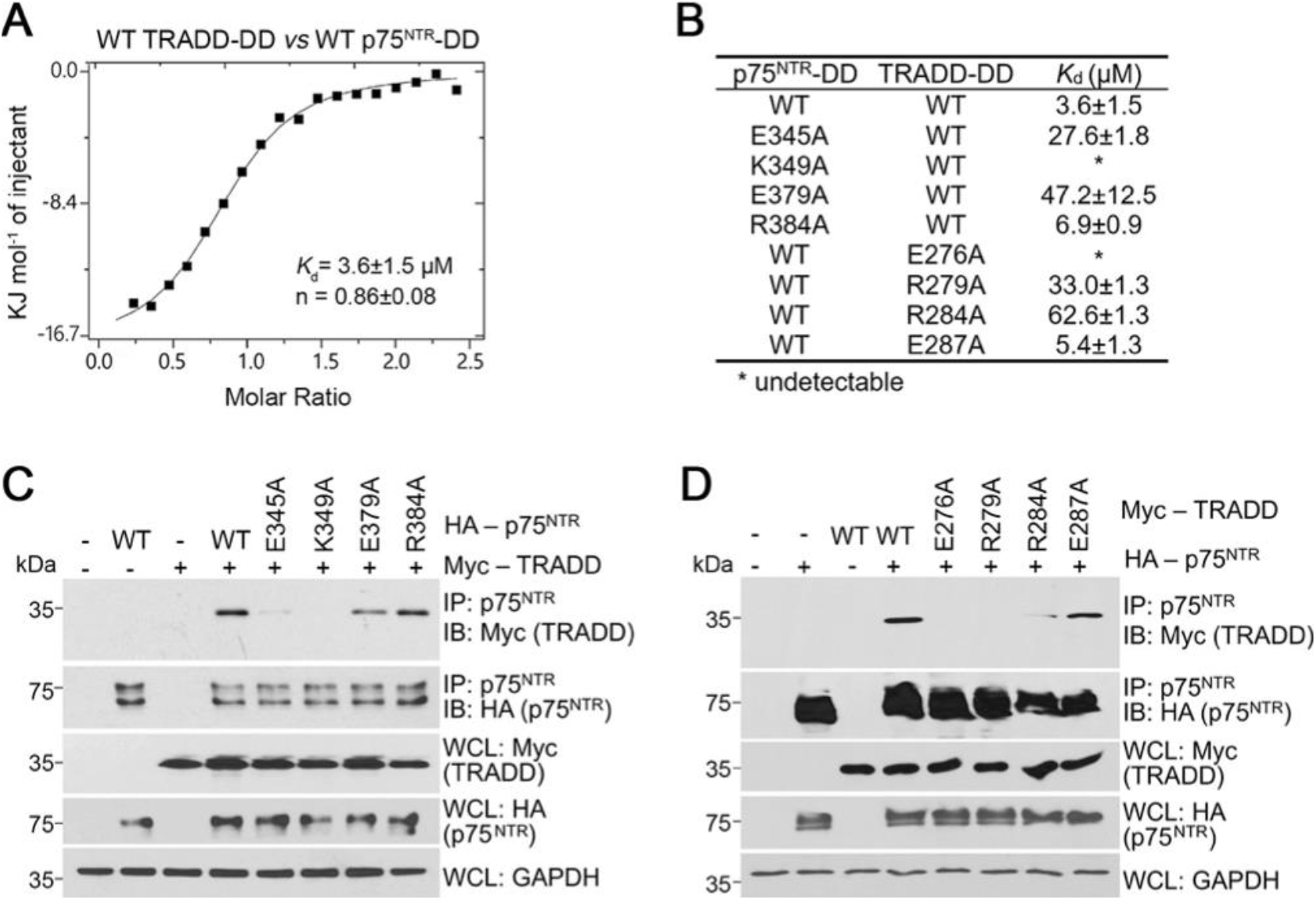
Mutagenesis studies. A) ITC binding curves of human TRADD to human p75NTR. B) Binding affinities expressed as dissociation constants (*K*d) of WT and point mutants of p75^NTR^ and TRADD derived from ITC data. C) Co-immunoprecipitation of wild type (WT) and point mutants of HA-tagged human p75^NTR^ with Myc-tagged human TRADD in transfected HEK 293T cells. IB, immunoblotting; IP, immunoprecipitation; WCL, whole cell lysate. The immunoblots shown are representative of three independent experiments. D) Co-immunoprecipitation of WT and point mutants of Myc-tagged human TRADD with HA-tagged human p75^NTR^ in transfected HEK 293T cells. The immunoblots shown are representative of three independent experiments.

### 2.2 p75^NTR^-DD interaction specificity

In p75^NTR^-mediated signaling pathways, the p75^NTR^-DD functions as a central hub for recruitment of various intracellular interactors or domains, including RhoGDI (Rho guanine nucleotide dissociation inhibitor), RIP2 (Receptor-interacting protein 2) kinase, TRADD, p45-DD, et al., in different cellular contexts. An important question is what could confer the interaction specificity of the p75^NTR^-DD. Although the p75^NTR^-DD exhibits a similar structural fold or 3D topology to other members in the DD superfamily, it has distinctive lengths and orientations of six α- and one 3_10_ helices, leading to unique surface features, such as protrusion or concave surfaces with different charge distribution, which are crucial for determining binding specificity of the p75^NTR^-DD (Figure 3).^[11b]^ Structural investigation of the complexes between the p75^NTR^-DD and its interactors reveals that the p75^NTR^-DD takes advantage of specific surfaces for recruiting different interactors (Figure 3). Mutagenesis studies on the charged or hydrophobic residues involved in the binding interfaces further provide functional validation of the structural insights obtained on the specific recognition of different interactors by the p75^NTR^-DD (Figure 2).^[16]^

**Figure 3.**
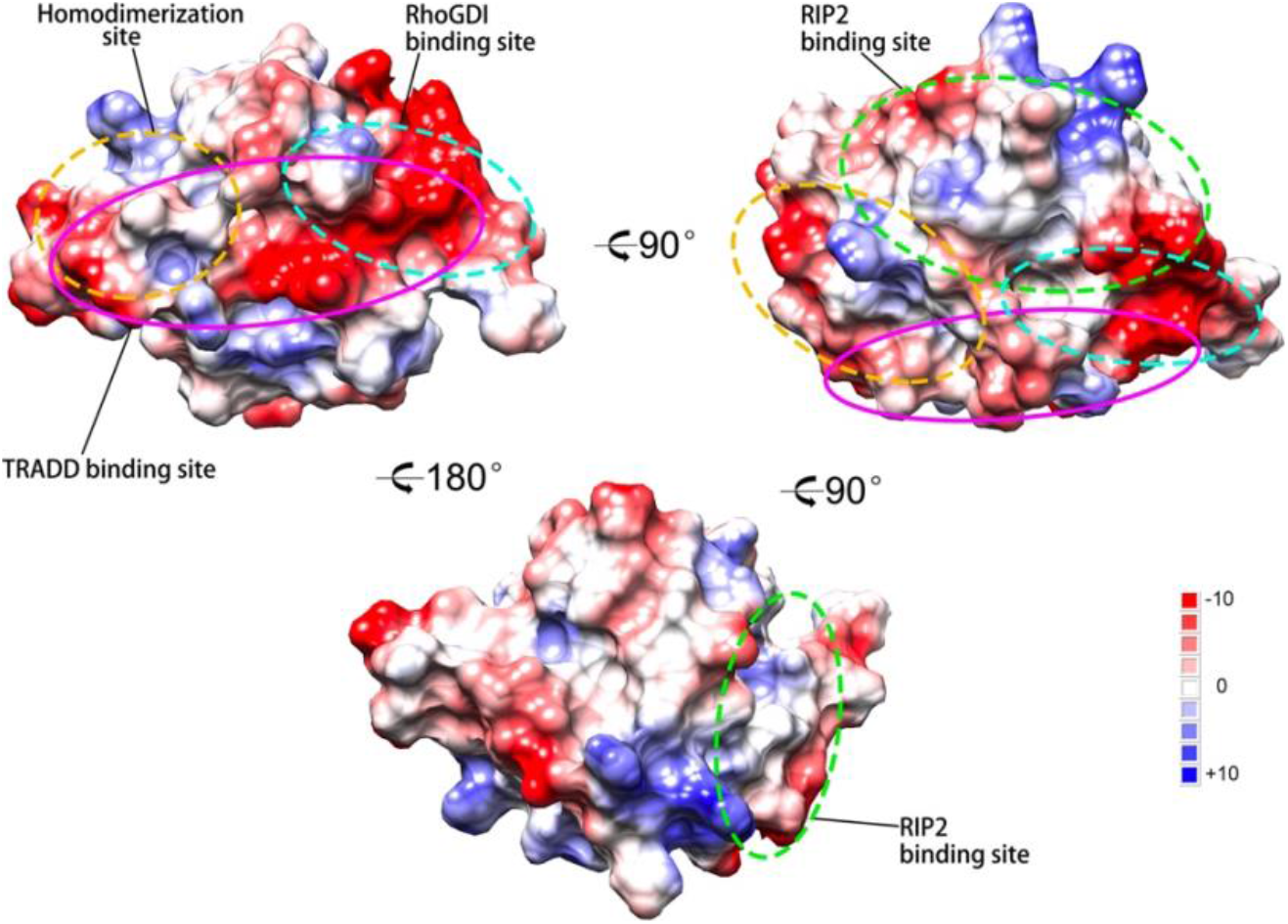
Charged surface of the p75^NTR^-DD and the binding sites for intracellular interactors. Unstructured regions are not shown. Color code is blue for positive charges, red for negative charges, and white for neutral surface. The patches on the surface of the p75^NTR^-DD responsible for binding TRADD, RhoGDI, RIP2 and the p75^NTR^-DD itself are circled in pink, cyan, green and brown, respectively.

The interaction specificity of the p75^NTR^-DD is not compromised by overlapping binding sites. Partially overlapping binding sites on the p75^NTR^-DD can lead to competitive binding of two or more proteins to the p75 ^NTR^-DD if they are co-expressed in a cell context. For example, the TRADD-DD binding site on the p75^NTR^-DD partially overlaps with p75^NTR^-DD homodimerization and RhoGDI binding sites (Figure 3). Therefore, recruitment of TRADD to p75^NTR^ requires separation of p75^NTR^-DD homodimer. Although co-expression of RhoGDI and TRADD in the same neuron cell is unknown, it is conceivable that signaling pathways regulated by RhoGDI and TRADD could not be activated at the same time.

### 2.3 Expression of TRADD in developing cerebellum

For TRADD to have a physiological role in p75^NTR^ signaling, it needs to be present in some of the same cells that express the receptor. We and others have previously reported on the expression and function of p75^NTR^ in developing cerebellar granule neurons (CGNs), the most abundant neuron subtype in the mammalian brain.^[9b, 14]^ In contrast, the specific expression of TRADD in distinct neuronal subpopulations, including CGNs, is not well understood. We therefore assessed whether TRADD is expressed in developing CGNs at early postnatal stages, coincident with the height of p75^NTR^ expression. Immunohistochemical analysis revealed that TRADD and p75^NTR^ were highly co-expressed in proliferating CGN precursors in the external granule layer (EGL) at postnatal day 2 (P2) (Figure 4A). Their expression levels were significantly lower at P5 (Figure 4B). By P7, the EGL is reduced, as CGNs mature and migrate to the internal granule layer (IGL), and both proteins were expressed at very low levels (Figure 4C). Thus, the spatiotemporal overlap of p75^NTR^ and TRADD expression is compatible with TRADD being involved in p75^NTR^ signaling during early stages of CGN development. Moreover, their predominant co-expression in early, proliferating CGN precursors suggest roles in cell-cycle progression and/or withdrawal.

**Figure 4.**
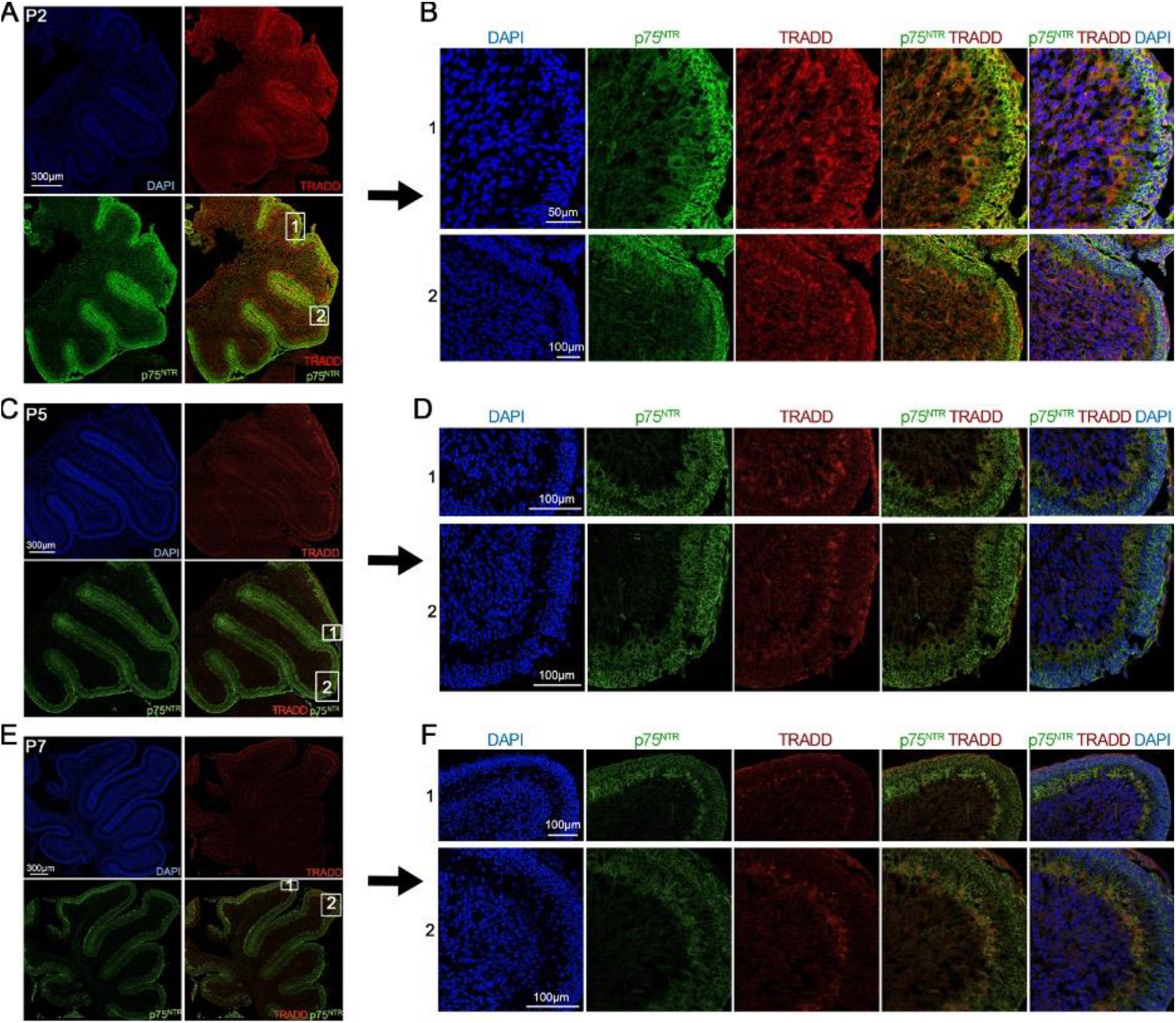
Expression of TRADD in developing cerebellum. Micrographs of a representative mid-sagittal section through the developing cerebellum of P2 (A/B), P5 (C/D) and P7 (E/F) wild type mouse stained with anti-TRADD (red) and anti-p75^NTR^ (green) antibodies, and counterstained with DAPI (blue). Higher magnification panels of P2, P5, and P7 are shown in (B), (D) and (F), respectively. Scale bars: (A), 300 μm; (B), 50 μm (upper) and 100 μm (lower); (C), 300 μm; (D), 100 μm; (E) 300 μm; (F), 100 μm.

### 2.4 Physiological relevance of TRADD/p75^NTR^ interaction in NF-κB signaling

It has been reported that interaction of TRADD with TNFR1 leads to activation of the NF-κB signaling pathway.^[17]^ In order to elucidate the functional relevance of the interaction between p75^NTR^ and TRADD, we investigated the activation state of this pathway in cultured CGNs derived from p75^NTR^ null mice after reconstitution with wild type p75^NTR^, the p75^NTR^ mutants K349A that is unable to bind TRADD or the p75^NTR^ mutant R384A that has a decreased interaction with TRADD. Our previous studies indicated that p75^NTR^ mediates tonic activation of NF-κB in cultured CGN neurons through endogenous production of NGF and autocrine stimulation of p75^NTR^ in these cells.^[9b, 14a]^ We assessed NF-κB activation by quantification of the levels of the p65 NF-κB subunit in the nuclei of cultured CGNs, a key step in canonical NF-kB signaling. In agreement with our previous studies, we observed reduced levels of p65 NF-κB in the nuclei of CGNs derived from p75^NTR^ knock-out mice, compared to wild type controls (Figure 5A,B). Importantly, transfection of knock-out CGNs with wild type p75^NTR^, but not K349A mutant, restored the levels of nuclear p65 NF-κB (Figure 5B). R384A mutant only partially restored NF-kB activation due to its weaker binding affinity to TRADD. This assessment indicates that interaction of TRADD with p75^NTR^ is required for normal activation of the NF-κB pathway in CGNs. Proximity ligation assay (PLA) further shows obvious complex formation between TRADD and p75^NTR^, while PLA signal is not detectable in KO CGN cells without transfected p75^NTR^ (Figure S5, Supporting Information).

**Figure 5.**
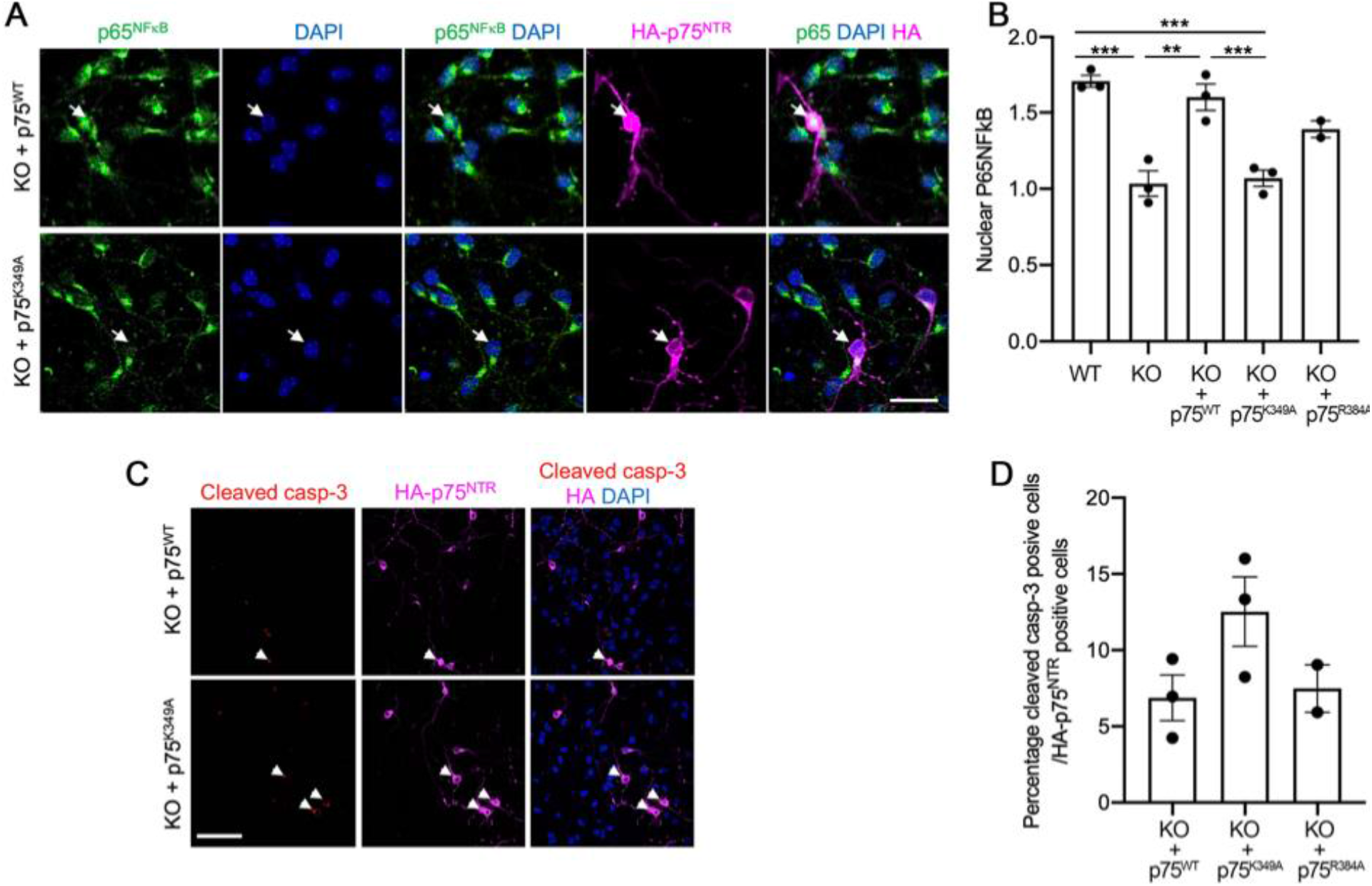
Functional role of TRADD/p75^NTR^ interaction in NF-κB signaling A) Representative micrographs of p75^NTR^ knock-out (KO) CGNs transfected with expression plasmids containing HA-tagged p75^WT^ (wild type) and HA-tagged p75^K349A^ mutant. After 2 days *in vitro,* cultures were immunostained with anti-P65 NF-κB (green) and anti-HA (magenta) antibodies, and counterstained with DAPI (blue). Transfected cells expressing HA constructs are indicated with white arrows. Scale bar, 20 μm. B) Quantification of nuclear p65 NF-κB (expressed as nuclear p65 fluorescence intensity /cytoplasmic p65 fluorescence intensity) in p75^NTR^ wildtype (WT) and knock-out (KO) CGNs transfected with HA-p75^WT^and HA-p75^K349A^ plasmids as indicated. Results are expressed as mean ± SEM from 3 independent cultures (*p< 0.05 and **p<0.01, one-way ANOVA followed by Turkey’s multiple comparison test). C) Representative micrographs of p75^NTR^ KO CGNs transfected with expression plasmids containing HA-tagged p75^WT^ (wild type) and HA-tagged p75^K349A^ mutant. After 2 days *in vitro,* cultures were immunostained with anti-caspase 3 (red), anti-HA (magenta) antibodies, and counterstained with DAPI (blue). Scale bar, 20 μm. D) Quantification of percentage cleaved caspase 3 positive cells (relative to DAPI) in p75^NTR^ KO CGNs transfected with HA-p75^WT^ and HA-p7^K349A^ plasmids as indicated. Mean ± SEM of densitometry from 3 separate cultures is shown (*p<0.05 and **p<0.01, one-way ANOVA followed by Turkey’s multiple comparison test).

As NF-κB activity has been shown to be essential for survival of CGNs,^[9b, 14a]^ we assessed apoptotic cell death in cultured CGNs from p75^NTR^ knock-out mice after reconstitution with wild type, K349A or R384A mutant by immunostaining for cleaved caspase-3. We found twice as many apoptotic cells in cultures reconstituted with the mutant p75^NTR^ K349A construct, compared to wild type (Figure 5C,D), in agreement with the inability of the former to efficiently activate NF-κB. Together, our expression and functional studies indicate that TRADD binding to p75^NTR^ is essential for the ability of the receptor to regulate NF-κB signaling and the balance between cell survival and cell death in CGNs, and perhaps other neuronal subpopulations and cell types outside the nervous system.

Although regulation of the NF-κB pathway by p75^NTR^ is often studied in the context of cell survival, there is also strong evidence for the importance of NF-κB signaling in cell cycle regulation, primarily through its of effects on the expression of several key components of the cell cycle machinery, including cyclin D1.^[18]^ Interestingly, recent work has indicated a role for p75^NTR^ signaling in the regulation of cell cycle duration and withdrawal in CGN precursors.^[14c, 19]^ Although those studies highlighted the role of the RhoA pathway in the effects of p75^NTR^ on the cell cycle, the NF-κB and RhoA signaling pathways are known to intersect on p75^NTR^, through competition between RhoGDI and RIP2 effectors for binding to the receptor DD.^[16a]^ Thus, it is possible that the interaction of TRADD with the p75^NTR^-DD described here plays a dual role in the early development of CGNs, both through direct regulation of the NF-κB pathway, as well as indirectly modulating RhoA signaling through competition for access to binding determinants in the DD.

### 2.5 Model for the initial stage of NF-κB signaling engaged by p75^NTR^ and TRADD

Based on the present studies, we propose a structural model for the initial stage of p75^NTR^ engagement with the TRADD/NF-κB pathway in developing cerebellar neurons (Figure 6). p75^NTR^ can form a disulfide-bound dimer at the plasma membrane, and homodimerization of two p75^NTR^-DDs closes their potential binding sites for downstream effectors. Since the N-terminal domain of TRADD (TRADD-NTD) can interact with the TRADD-DD,^[20]^ TRADD could also exist in a closed state before recruitment to the receptor although the exact binding interface between the TRADD-NTD and the TRADD-DD is unclear. p75^NTR^ signaling through TRADD depends on NGF, which is expressed by developing cerebellar neurons. NGF binding to the p75^NTR^-ECD leads to separation of intracellular p75^NTR^-DDs through disulfide-bonded p75^NTR^-TMD.^[16a]^ Since p75^NTR^-DD homodimerization site partially overlaps with TRADD-DD binding site on the p75^NTR^-DD (Figure 3 and Figure S6, Supporting Information), dissociation of p75^NTR^-DD homodimer is required for the recruitment of the TRADD-DD to p75^NTR^, which could lead to the opening of two domains of TRADD. The free TRADD-NTD is envisioned to further recruit as yet unknown downstream molecules to activate the NF-κB pathway and promote neuron cell survival. In TNFR1- and DR3-mediated signaling pathways, the three molecules of TRADD-NTD were shown to associate with trimeric TRAF domains of TNF receptor-associated factor 2 (TRAF2) to initiate NF-κB signaling.^[11a, 21]^ Since the unstructured linker between the TRADD-NTD and the TRADD-DD is long (~37 amino acid residues), it is possible that three p75^NTR^ dimers are clustered together by two TRAF2 trimers on the membrane through TRADD and form a larger complex if TRAF2 is co-expressed with TRADD and p75^NTR^ in developing CGNs. In this scenario, opening of two-domain TRADD is also necessary since key residues of the TRADD-NTD for binding the TRADD-DD are involved in the interaction with TRAF2 (Figure S7, Supporting Information).^[20]–[21]^

**Figure 6.**
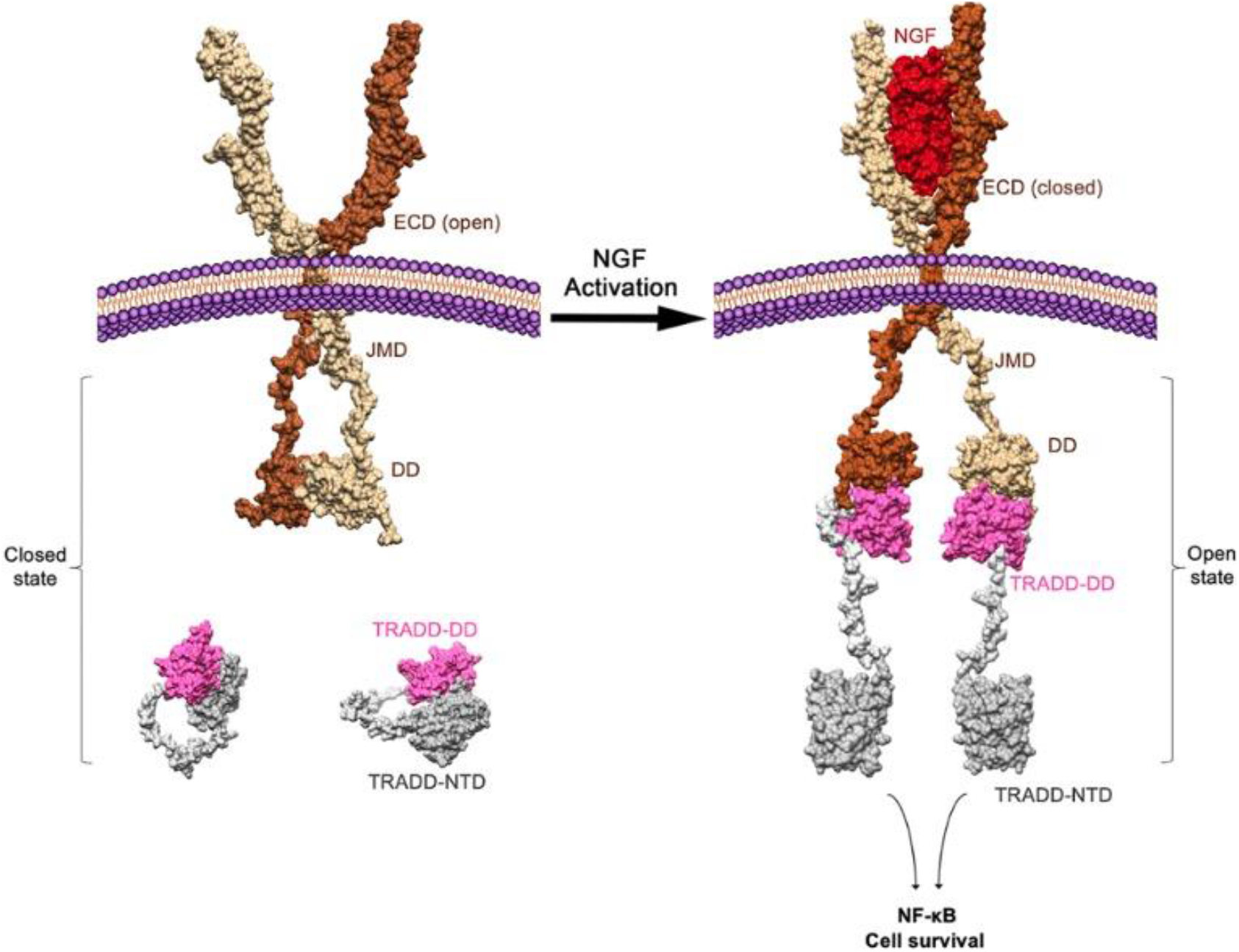
Structure model of p75^NTR^ NF-κB signaling via the recruitment of TRADD. p75^NTR^ model was built based on available structures of individual domains and domain complexes. The domain orientation and interface between TRADD-NTD and TRADD-DD are not defined in this cartoon.

Previous studies have demonstrated that the caspase recruitment domain (CARD domain) of RIP2 interacts with the p75^NTR^-DD to activate NF-κB pathway in developing cerebellum as well as in Schwann cells.^[9a, 14a, 22]^ It is interesting to know that if RIP2 and TRADD could compete with each other to bind to the p75^NTR^-DD. Structural docking shows that the p75^NTR^-DD, the TRADD-DD, and the RIP2-CARD could form a tripartite complex due to non-overlapping binding interfaces (Figure S8, Supporting Information). Co-immunoprecipitation experiments performed in HEK 293T cells further demonstrated that mutations of key residues at the binding interface of p75^NTR^-DD:TRADD-DD complex do not reduce the capacity of p75^NTR^ to bind RIP2 (Figure S9, Supporting Information). Therefore, it is possible that p75^NTR^ recruits both TRADD and RIP2 upon NGF binding for downstream NF-κB signaling in developing CGNs if RIP2 is co-expressed with TRADD. RIP2 was also shown to diminish the interaction between TRAF6 and p75^NTR^ through a steric hindrance effect on the receptor itself. Nevertheless, whether TRADD binding to the death domain of p75^NTR^ interferes with TRAF6 binding remains to be investigated. Given that neither TRADD binds to the same residues of p75^NTR^ as TRAF6 does and the MW of RIP2 (≈60kDa) is nearly twice as big as TRADD (≈34kDa), TRADD could be too small to interfere with TRAF6 binding in the same way RIP2 is capable of.

## 3. Conclusion

In summary, here we report the structural mechanism underlying recruitment of TRADD to p75^NTR^ through DD interactions and reveal that interaction specificity of the p75^NTR^-DD, an intracellular signaling hub, relies on its distinct surface patches. We also identify the crucial role of TRADD in p75^NTR^-mediated NF-κB signaling and cell survival in CGNs. Our results provide a new structural framework for understanding the mechanisms by which this adaptor protein creates a large complex on p75^NTR^ to propagate downstream signaling.

## 4. Experimental Section

### Protein Expression and Purification

The cDNAs of human p75^NTR^ DD (330-427) and TRADD-DD (199-312) were amplified from total human embryonic stem (ES) cell cDNA and subcloned into pET32-derived expression vectors between BamH I and Xho I restriction sites. Unlabeled proteins were expressed in *E.coli* strain SoluBL21 (DE3) in M9 medium. All protein samples were purified by Ni-NTA affinity chromatography, ionic exchange (MonoQ or MonoS) and/or gel filtration (Superdex 75). Isotopic labelling of ^15^N and/or ^13^C was carried out by expressing the proteins in M9 minimal medium containing 15N-NH_4_Cl and/or 13C-labeled glucose as the sole source of nitrogen and carbon. For protein complexes, two double-labelled samples were prepared in pure water with 10 × 10−3 М D_10_-DTT: (1) 0.8 × 10−3 М 13C, ^15^N-labeled p75^NTR^ DD mixed with 1.0 × 10^−3^ М unlabeled TRADD-DD; (2) 0.8 × 10^−3^ М ^13^C, ^15^N-labeled TRADD-DD mixed with 1.0 × 10^−3^ М unlabeled p75^NTR^ DD.

### NMR Spectroscopy Experiments and Structure Determination

All NMR experiments were carried out on a Bruker 800 MHz NMR spectrometer (AVANCE) with a cryogenic probe at 301K. All the NMR spectra were processed with NMRPipe ^[23]^ and analyzed with NMRDraw and NMRView supported by a NOE assignment plugin.^[24]^ Sequence-specific assignments of backbone and side chains were obtained by using previously described methods.^[25]^ The chemical shift values of backbone Cα and Cβ were analyzed by TALOS+ to predict backbone dihedral angles.^[26]^ Intramolecular NOEs were assigned from 4D time-shared ^13^C, ^15^N-edited NOESY spectra.^[27]^ Ambiguous NOEs were obtained with iterated structure calculations by CYANA.^[28]^ Based on NOE peak volume, NOE values were binned into short (1.8-2.8 Å), medium (1.8-3.4 Å) and long (1.8-5.5 Å) distances. Final structure calculation was started from 100 conformers. 10 conformers with the lowest target function values were selected for energy minimization in AMBER force field.^[29]^ PROCHECK-NMR was used to assess the quality of the structures.^[30]^ All the structural figures including charged surfaces were made using UCSF Chimera.^[31]^

### Model structure calculations

The structure of p75^NTR^-DD:TRADD-DD:RIP2-CARD complex was modeled using HADDOCK software.^[32]^ The lowest-energy monomeric conformers of p75^NTR^-DD, TRADD-DD and RIP2-CARD were used as starting structures. Key residues for domain interactions identified in p75^NTR^-DD:TRADD and p75^NTR^-DD:RIP2-CARD complexes were selected as active residues, and surface-exposed residues within a 6.5 Å radius around the active residues were used as passive ones. 1,000 rigid-body docking structures were generated by energy minimization. The best 100 structures with lowest intermolecular energies were selected for semi-flexible simulated annealing in torsion angle space followed by a final refinement in explicit water. The final structure with the lowest HADDOCK score was used to represent the structure model.

### ITC binding assay

All protein samples used in the ITC binding assay were prepared in deionized water with 1.4 × 10^−3^ М β-mercaptoethanol. ITC binding assay was performed via an ITC200 (GE Healthcare) equipment at 25 °C. To determine the binding affinity, 0.4 × 10^−3^ М WT TRADD-DD was titrated into 0.04 × 10^−3^ М WT p75^NTR^-DD. Mutagenesis studies were performed in a similar way. The thermograms were integrated by the Origin software and fitted to single-site binding model. The standard deviation of each *K*d value was calculated from three independent ITC titration experiments.

### Animals

Mice were housed in a 12-hour light/dark cycle and fed a standard chow diet. The transgenic mouse lines used were p75^NTR^ knockout (KO).^[33]^ Transgenic mice were maintained in a c57bl6J background. Mice of both sexes were used for the experiments. All animal experiments were conducted in accordance with the National University of Singapore Institutional Animal Care. The lab certificate/approval number is OSHM/PI/14/SOM-250.

### Plasmids

Full-length HA-tagged p75^NTR^ and full-length Myc-tag TRADD were expressed from a pcDNA3 vector backbone (Invitrogen). The HA-p75^E345A^, HA-p75^K349A^, HA-p75^E379A^, HA-p75^R384A^, Myc-TRADD^E276A^, Myc-TRADD^R279A^, Myc-TRADD^R284A^ and Myc-TRADD^E287A^ mutant constructs were selected based on p75^NTR^-DD:TRADD-DD complex structure.

### Neuronal cultures

Wild type and p75^NTR^ KO CGNs were trypsinized and plated at a density of 40 000 (for cell death assay) or 200 000 (for protein collection) cells per coverslip coated with poly-L-lysine (Sigma, Cat: P7280) in a 24-well (Starlab) in neuro basal medium supplemented with B27 (Gibco, Cat: 17504001, 25 × 10^−3^ М KCl (Sigma, Cat: P9541), 1× 10^−3^ М glutamine (Gibco, Cat:25030149) and 2 mg/ml gentamicin (Invitrogen, Cat: 15750060).

### Transfection

Neurons were transfected with either HA-p75^WT^, HA-p75^K349A^ or HA-p75^R384A^ plasmids using Neon™ transfection system (Thermo scientific, Cat: MPK10025) prior to plating. 250ng plasmid per well in the 24-well plate was used.

### Cell death assay

For assessing apoptosis, p75^NTR^ KO neurons transfected with the different plasmids (see above) were cultured for 2 days, fixed with solution containing 4% paraformaldehyde and 4% sucrose. The fixed cells were labeled with cleaved caspase 3 (Cell signal, 9761, 1:400), anti HA-tag (Invitrogen, 71-5500, 1:250) and DAPI. For each experiment, neurons were culture in duplicates and at least 15 images were taken per coverslip.

### Proximity ligation assay (PLA)

After transfection or treatment, CGNs were fixed for 15 min in 4% paraformaldehyde (PFA)/4% sucrose, permeabilized, and blocked in 10% normal donkey serum and 0.3% Triton X-100 in PBS. Cells were then incubated overnight at 4 °C with anti-p75^NTR^ (Promega; G323A; 1:300) and anti-TRADD (Santa Cruz; sc-46653; 1:500) antibodies in PBS supplemented with 3% BSA. The Duolink In Situ Proximity Ligation kit (Sigma; DUO92007) was used as per the manufacturer’s instructions with fluorophore-conjugated secondary antibody to recognize HA (Sigma; H3667, 1:250) included during the amplification step. Transfected and HA-positive cells were imaged with an LSM Imager Z2 confocal microscope (Zeiss) to detect PLA signals.

### Immunohistochemistry

P2, P5 and P7 wild type animals were perfused first with PBS, followed by 4% paraformaldehyde. Harvested cerebellar were post fixed in 4% paraformaldehyde for 16 h and cryoprotected in 30% sucrose before freezing. OCT-embedded cerebellar were frozen at −80°C overnight and serially sectioned at 30 μm in the sagittal plane using cryostat. Midline sections were mounted onto electrostatic charged slides (Leica Microsystems), blocked with 5% donkey serum (Fisher scientific) containing 0.3% Triton X-100 (Sigma) in PBS for 1 h at room temperature and then incubated for 16 h at 4°C with primary antibodies. The sections were washed in PBS before incubated with the appropriate secondary antibodies.

### Immunocytochemistry

the cultures were fixed in 4% paraformaldehyde and 4% sucrose for 15 min and washed with PBS before blocking nonspecific binding and permeabilizing with blocking solution (5% donkey serum and 0.3% Triton X-100) in PBS for 1 h at room temperature. Neurons were incubated overnight with the primary antibodies in 1% blocking solution at 4°C. After washing with PBS, the cultures were incubated with the appropriate secondary antibodies. The primary antibodies used in this study were: polyclonal anti-p75^NTR^ (Neuromics, GT15057, 1:250), monoclonal anti-TRADD (Santa Cruz, sc-46653, 1:500), polyclonal anti-HA (Sigma, H3667, 1:250), polyclonal anti-P65 NF-κB (Santa cruz, sc-372, 1:500) and polyclonal anti-cleaved caspase 3 (Cell signal, 9761, 1:400), Secondary antibodies were Alexa Fluor–conjugated anti-immunoglobulin from Life Technologies, Invitrogen, used at 1:1000 (donkey anti-rabbit IgG Alexa Fluor 555, A31572, donkey anti-goat IgG Alexa Fluor 488, A11055, donkey anti-mouse IgG Alexa Fluor 555, A31570 and donkey anti-mouse IgG Alexa Fluor 647, A31571). Images were obtained using a Zeiss Axioplan confocal microscope.

### Immunoprecipitation

HEK 293T cells were transfected with the polyethylenimine (PEI) method. 48 hours post transfection, cells were lysed in lysis buffer (50 × 10^−3^ М Tris/HCl pH 7.5, 1 × 10^−3^ М EDTA, 270 × 10^−3^ М Sucrose, 1% (v/v) Triton X-100, 0.1% (v/v) 2-mercaptoethanol and 60 × 10^−3^ М n-Octyl-β-D-Glucopyranoside) containing protease inhibitor (Roche). Total protein was collected and incubated with anti-p75^NTR^ (Alomone, ANT-007, 1ug) overnight at 4 °C and then incubated with Sepharose protein-G beads (GE Healthcare). Samples were then prepared for immunoblotting as described below.

### Immunoblotting

Immunoblotting protein samples were prepared for SDS-PAGE in SDS sample buffer (Life Technologies) and boiled at 95 °C for 10 min before electrophoresis on 12% gels. Proteins were transferred to PVDF membranes (Amersham). Membranes were blocked with 5% non-fat milk and incubated with primary antibodies. The following primary antibodies were used at the indicated dilutions: mouse anti-Myc (Roche, 1:3000), goat anti-p75^NTR^ (Neuromics, GT15057, 1:3000) and anti-GAPDH (Sigma, G9545, 1:15000). Immunoreactivity was visualized using appropriate HRP-conjugated secondary antibodies. Immunoblots were developed using the ECL Advance Western blotting detection kit (Life Technologies) and exposed to Kodak X-Omat AR films.

### Statistical analysis

Data are expressed as mean and standard errors. No statistical methods were used to predetermine sample sizes but our sample sizes are similar to those generally used in the field. Following normality test and homogeneity variance (Kolmogorov-Smirnov test or Brown-Forsythe test), group comparison was made using one-way ANOVA followed by Turkey’s post-hoc test. Differences were considered significant for *P* < 0.05. The experiments were not randomized. Data from all experiments are included; none were excluded.

## Supporting Information

Supporting Information is available from the Wiley Online Library or from the author.

## Acknowledgements

We thank Dr Jing-song Fan for NMR data collection, and Miss Ket Yin Goh for technical assistance. This research was supported by funds from National Natural Science Foundation of China (Grant No. 21974093) to Z. Lin.; as well as Singapore National Medical Research Council (NMRC/CBRG/0107/2016), Singapore Ministry of Education (MOE2018-T2-1-129), and Swedish Research Council (VR-2016-01538) to C.F.Ibanez.

Received: ((will be filled in by the editorial staff))

Revised: ((will be filled in by the editorial staff))

Published online: ((will be filled in by the editorial staff))

