## Supporting Information for "Structural and functional elucidation of NF-κB signaling by the p75 neurotrophin receptor through recruitment of TRADD"

N. Zhang, Z. Li, Dr. W. Yuan, Prof. Z. Lin
Tianjin Key Laboratory of Function and Application of Biological Macromolecular Structures, School of Life Sciences, Tianjin University, Tianjin 300072, P.R. China

Dr. L. Kisiswa
Department of Biomedicine, Aarhus University, Aarhus DK-8000, Denmark

Prof. Z. Lin, Dr. A. Ramanujan, E. W. Sim, Prof. C. F. Ibáñez
Department of Physiology & Life Sciences Institute, National University of Singapore, 117456, Singapore

Prof. C. F. Ibáñez
Department of Neuroscience, Karolinska Institute, Stockholm 17165, Sweden

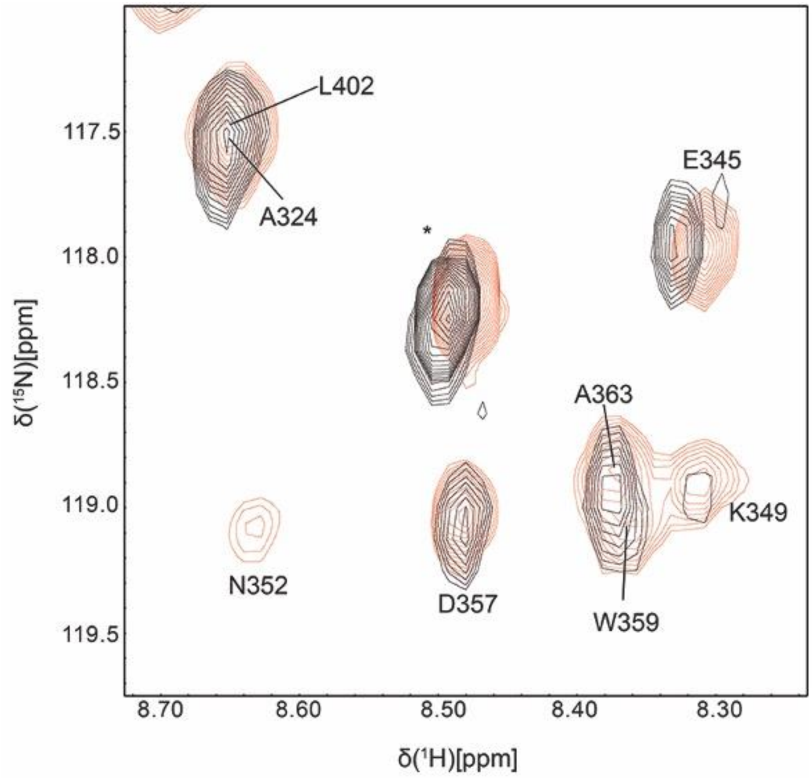


**Figure S1.** Expanded region of [$^{1}$H-$^{15}$N] HSQC spectra of the p75$^{NTR}$-DD in the absence (black) and presence (red) of the TRADD-DD at 28ºC in pure water. The concentration of the p75$^{NTR}$-DD is ~0.8 mM and the molar ratio of p75$^{NTR}$-DD to TRADD-DD is ~1:1.2. *: Ser residue from His tag. Upon binding the TRADD-DD, Intensities of cross peaks from residues K349 and N352 significantly increased.

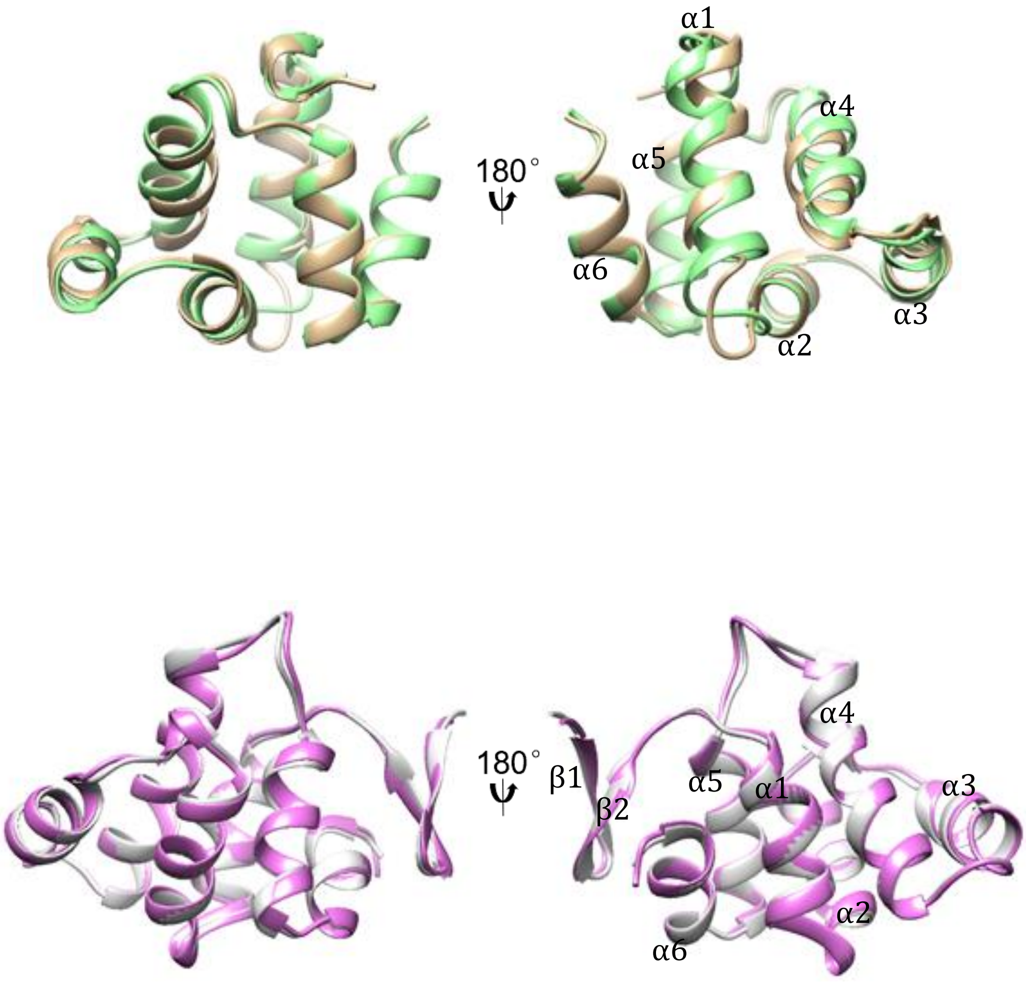


**Figure S2.** Structural comparisons. A) Structural overlap of p75$^{NTR}$-DD (brown) from p75$^{NTR}$-DD:TRADD-DD complex and from p75$^{NTR}$-DD homodimer (green). The backbone R.M.S.D. is ~1.1 Å. B) Structural overlap of TRADD-DD (pink) from p75$^{NTR}$-DD:TRADD-DD complex and from monomeric TRADD-DD (grey). The backbone R.M.S.D. is ~0.50 Å. The secondary structure elements are labeled.

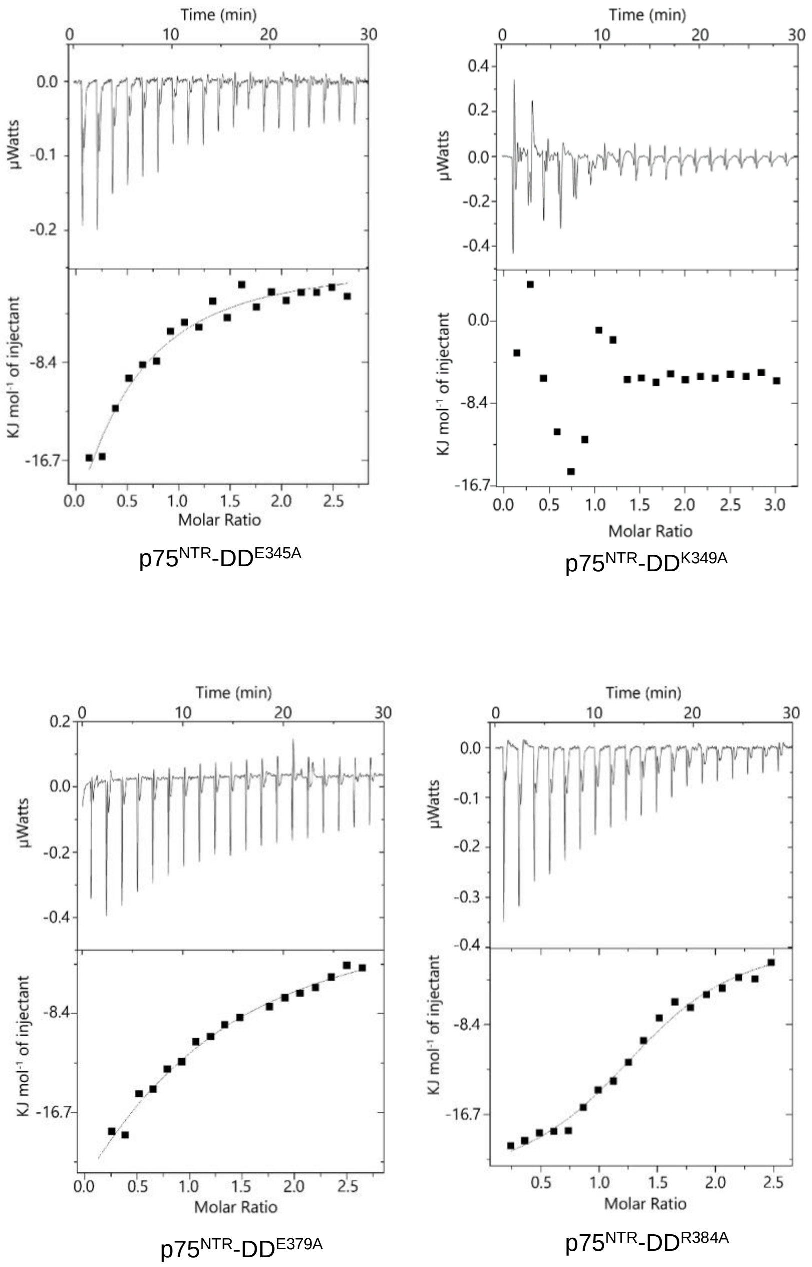


**Figure S3.** ITC binding curves of point mutants of the p75$^{NTR}$-DD to WT TRADD-DD.

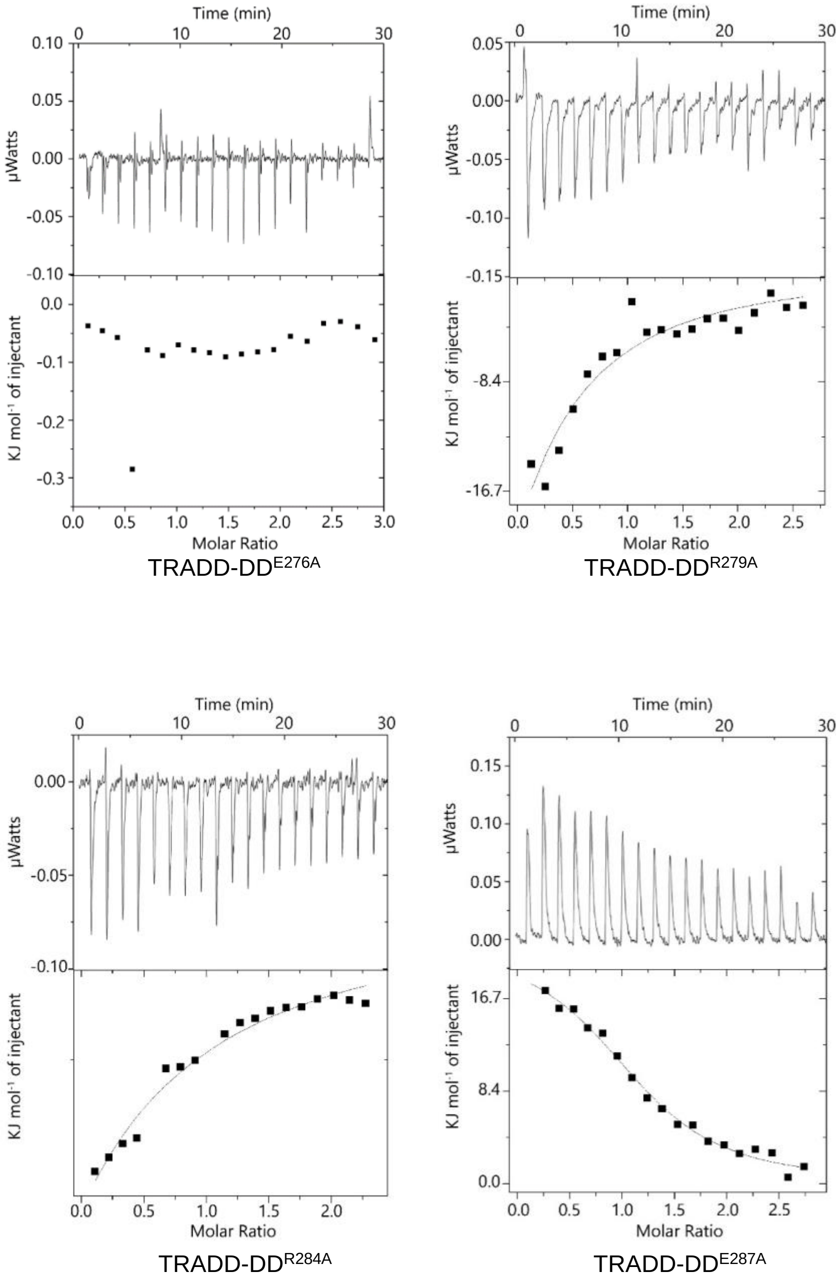


**Figure S4.** ITC binding curves of point mutants of the TRADD-DD to WT p75$^{NTR}$-DD.

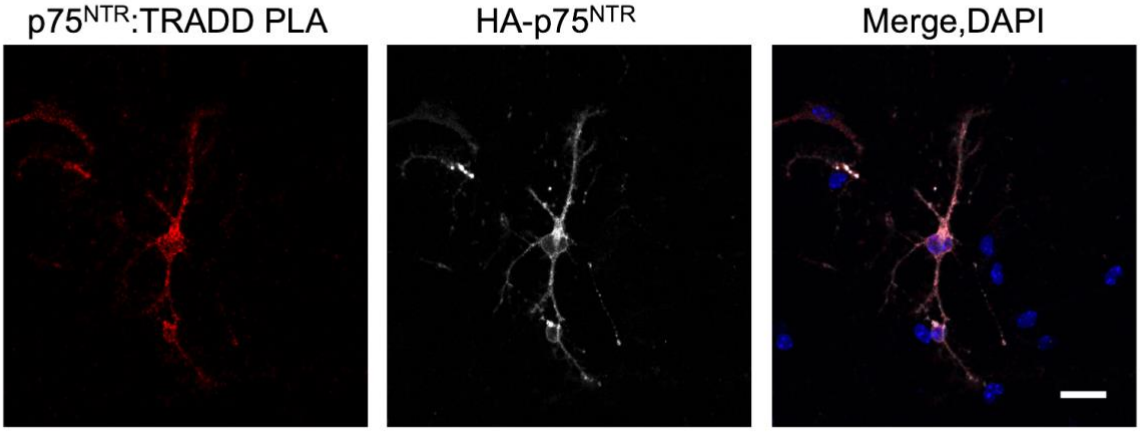


**Figure S5.** Interaction between p75$^{NTR}$ and TRADD in CGN

Micrographs of p75$^{NTR}$/TRADD PLA (red) in p75$^{NTR}$ knock-out transfected with expression plasmids containing HA-tagged WT p75$^{NTR}$ cultured for 2 days in vitro. Cells were counterstained with antibodies HA and DAPI. Scale bar, 20mM.

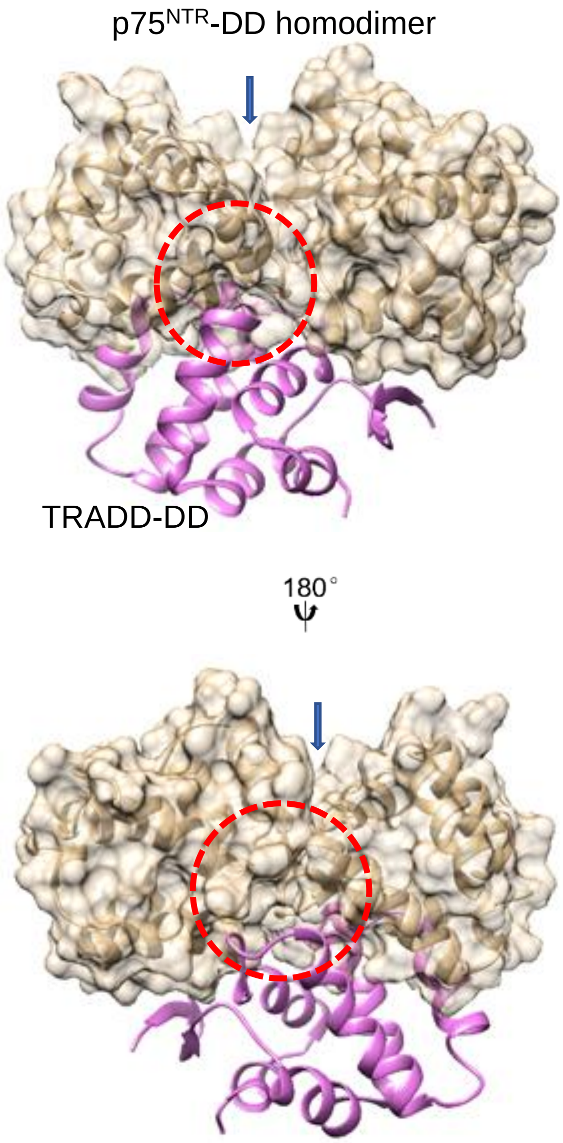


**Figure S6.** p75$^{NTR}$-DD homodimer interface partially overlaps with TRADD-DD binding site on p75$^{NTR}$-DD monomer. Surface representation of p75$^{NTR}$-DD homodimer (brown) with overlapped ribbon drawing of TRADD-DD (pink) demonstrates steric clash between TRADD-DD and one of p75$^{NTR}$-DD monomer. The homodimerization interface is indicated by arrow head. The partially overlapping interface are circled in red.

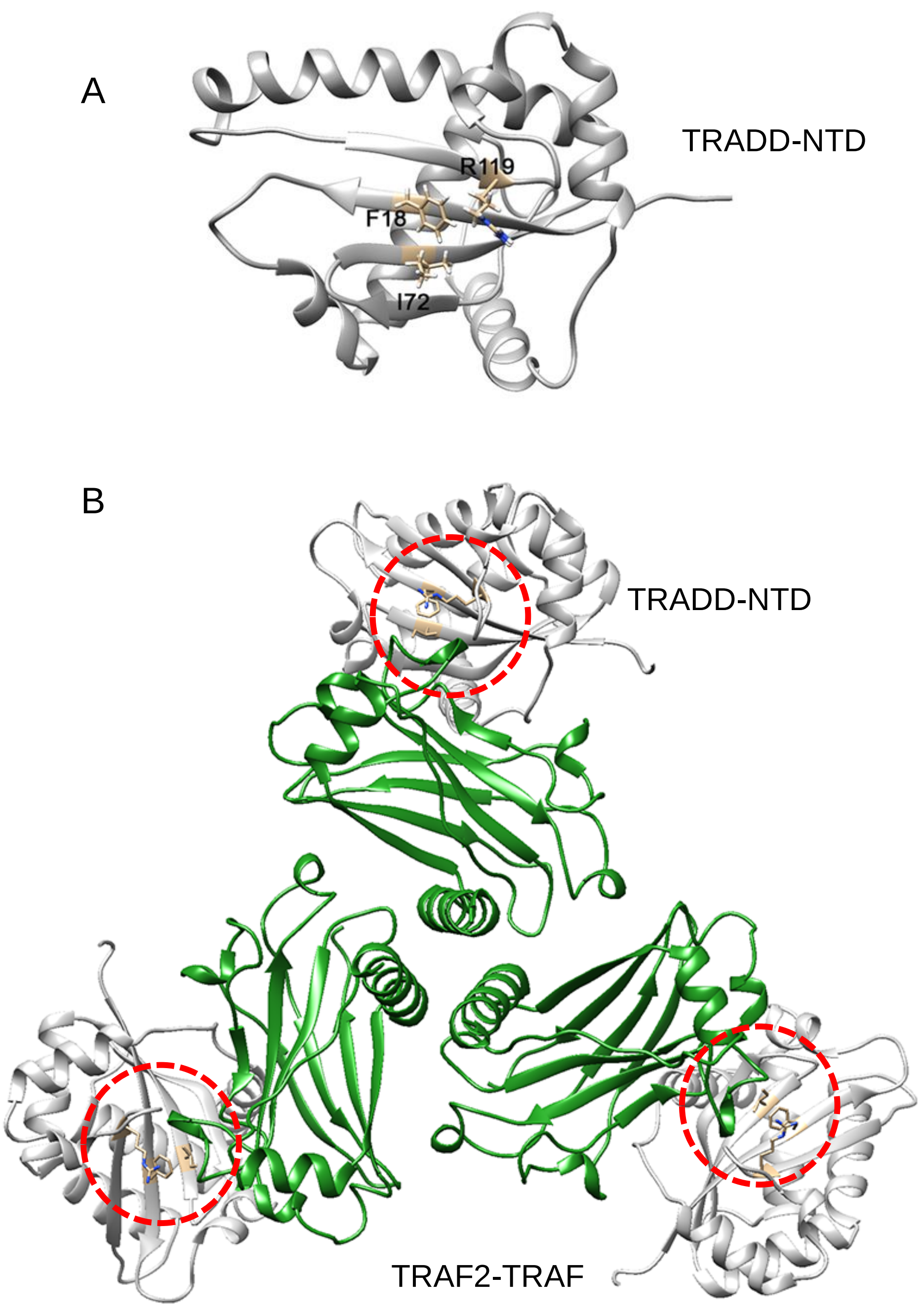


**Figure S7**. Binding of the TRADD-DD and TRAF2 to the TRADD-NTD at partially overlapping epitopes. A) Solution structure of monomeric TRADD-NTD (grey). PDB ID: 1F2H. Three key restudies (F18, I72, and R119) involved in binding the TRADD-DD are depicted in ball and stick. B) Crystal structure of complex between the TRADD-NTD (grey) and the TRAF domain of TRAF2 (green). PDB ID:1F3V. Residues F18, I72, and R119 from the TRADD-NTD are shown and circled in red.

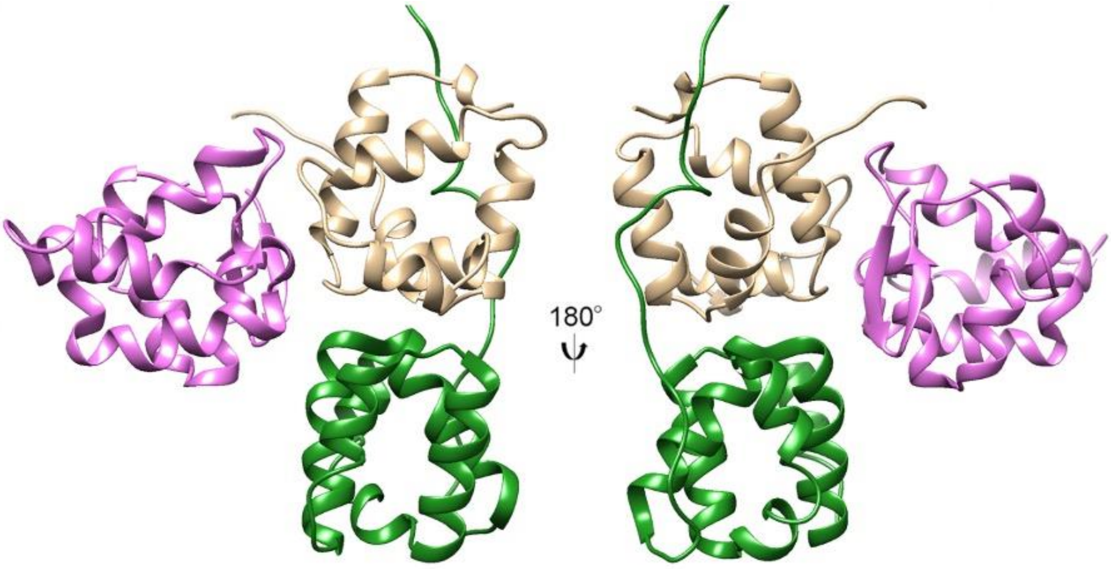


**Figure S8**. HADDOCK structure of the tripartite complex between the p75$^{NTR}$-DD (brown), the TRADD-DD (pink) and the RIP2-CARD (green).

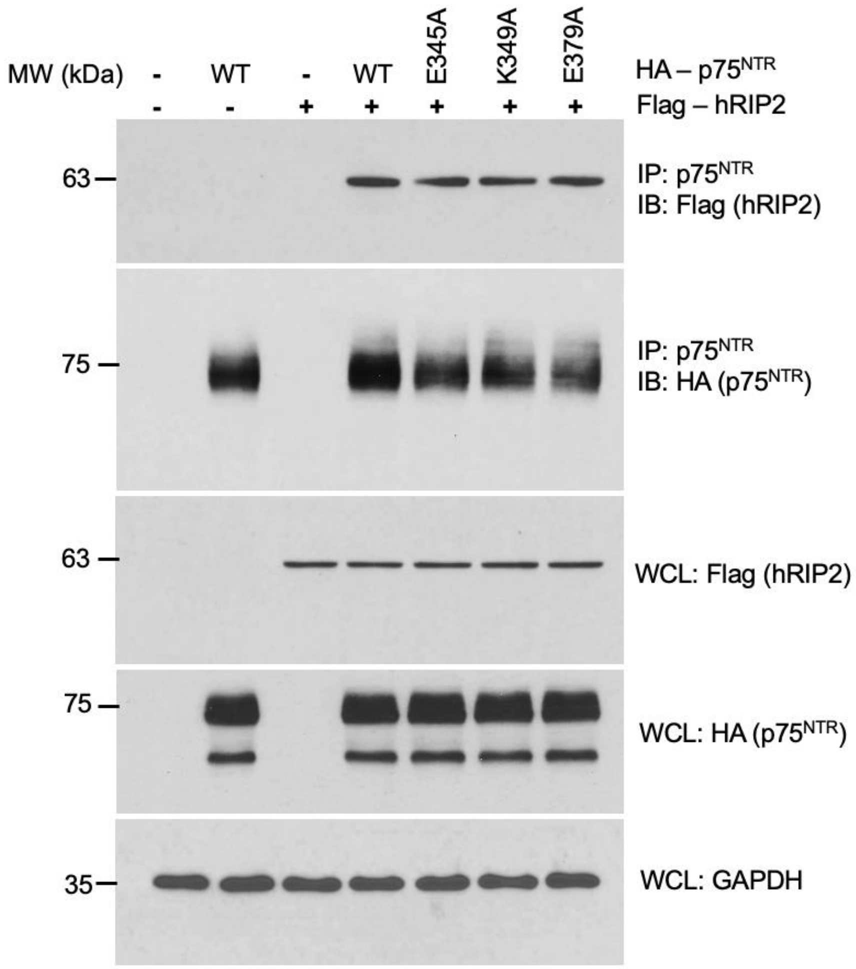


**Figure S9**. Co-immunoprecipitation of wild type (WT) and point mutants of HA-tagged human p75$^{NTR}$ with Flag-tagged human RIP2 in transfected HEK 293T cells. IB, immunoblotting; IP, immunoprecipitation; WCL, whole cell lysate.
